# Single-cell genomics of uncultured bacteria reveals dietary fiber responders in the mouse gut microbiota

**DOI:** 10.1101/784801

**Authors:** Rieka Chijiiwa, Masahito Hosokawa, Masato Kogawa, Yohei Nishikawa, Keigo Ide, Chikako Sakanashi, Kai Takahashi, Haruko Takeyama

## Abstract

**Background:** The gut microbiota can have dramatic effects on host metabolism; however, current genomic strategies for uncultured bacteria have several limitations that hinder their ability to identify responders to metabolic changes in the microbiota. In this study, we describe a novel single-cell genomic sequencing technique that can identify metabolic responders at the species level without the need for reference genomes, and apply this method to identify bacterial responders to an inulin-based diet in the mouse gut microbiota.

**Results:** Inulin feeding changed the mouse fecal microbiome composition to increase *Bacteroides* spp., resulting in the production of abundant succinate in the mouse intestine. Using our massively parallel single-cell genome sequencing technique, named SAG-gel platform, we obtained 346 single-amplified genomes (SAGs) from mouse gut microbes before and after dietary inulin supplementation. After quality control, the SAGs were classified as 267 bacteria, spanning two phyla, four classes, seven orders, and 14 families, and 31 different strains of SAGs were graded as high- and medium-quality draft genomes. From these, we have successfully obtained the genomes of the dominant inulin-responders, *Bacteroides* spp., and identified their polysaccharide utilization loci and their specific metabolic pathways for succinate production.

**Conclusions:** Our single-cell genomics approach generated a massive amount of SAGs, enabling a functional analysis of uncultured bacteria in the intestinal microbiome. This enabled us to estimate metabolic lineages involved in the bacterial fermentation of dietary fiber and metabolic outcomes such as short-chain fatty acid production in the intestinal environment based on the fibers ingested. The technique allows the in-depth isolation and characterization of uncultured bacteria with specific functions in the microbiota and could be exploited to improve human and animal health.

## Background

The gut microbiota plays a crucial role in the control of host physiology and metabolism through the release and transformation of metabolites like short-chain fatty acids (SCFAs) [1, 2]. Many studies have demonstrated that dietary fiber supplementation can modulate gut microbiota composition and promote SCFA production [3–5]. This dietary fiber metabolism is particularly important for non-digestible inulin-type fructans classified as prebiotics, which are substrates that promote the growth of beneficial microorganisms in the gut [6–12]. Previous reports indicated that several bacterial taxa in the intestine, including *Firmicutes, Bacteroides,* and *Bifidobacterium,* utilize fructan, and that dietary fructan can result in the growth of *Actinobacteria, Firmicutes,* or *Bacteroides.* However, our understanding of nutrient sensing and utilization by gut microbiota is limited, and there is a lack of predictability of the response of the microbiota to such dietary interventions. Thus, to enhance the benefits of dietary fiber to the host, it is important to identify inulin-responders in the complex microbiome community of the host intestine and characterize their mechanisms of inulin fermentation and potential for SCFA production.

Recently developed metagenomic approaches can provide comprehensive microbial profiling and putative functions using 16S rRNA gene amplicon sequencing and shotgun metagenome sequencing. These methods have enabled the exploration of complex communities in the gut environment without the need for *in vitro* cultivation of the intestinal microbiome. However, 16S rRNA gene sequencing provides snapshots of microbial composition, but does not provide information on microbial metabolic functions [13]. Shotgun metagenomics provides the complete genomic content of the microbial community, but it is generally difficult to link taxonomic and functional information for individual microbial strains [14]. Importantly, these techniques rely on the availability of reference microbial genomes for accurate functional assignments of specific organisms and precise taxonomic classification in the microbiome [15–17]. However, an estimated half of human gut species lack a reference genome. Thus, a new culture-independent genomic approach is required to enhance the reference genomes along with conventional metagenomic approaches and unveil the taxonomy and functions of the yet uncultured microbes at the strain level.

In this study, we used single-cell genome sequencing to identify microbial responders of dietary fiber, inulin, in mouse gut microbiota. Our massively parallel single bacterial genome sequencing technique, named single-cell amplified genomes in gel beads sequencing (SAG-gel), utilizes a gel matrix that enables a serial enzymatic reaction for tiny genome amplification and provides a support for the isolation of the amplified genomes. We targeted the family *Bacteroidaceae* from massively produced single-cell amplified genomes and characterized their polysaccharide utilization loci (PUL) and SCFA production pathways [7, 8, 11].

Near-complete genomes obtained via single-cell genome sequencing provided us new insights into the mechanisms of microbial proliferation and SCFA production in the intestine. The resulting genome set can serve as the basis for strain-specific comparative genomics to understand uncultured organisms in the gut microbiome. These findings will help us to predict the metabolic fermentation of dietary fibers based on the presence and ability of the specific responders.

## Results

### Effects of dietary fiber inulin on gut microbiota composition

First, we assessed changes in mouse gut microbiota composition before and after inulin supplementation. To account for diurnal oscillations in gut microbiota composition [18, 19], feces were sampled in the morning and evening after two weeks of continuous inulin feeding. Cellulose, which is generally poorly fermented by gut microbes, was used as a control diet fiber. As shown in Figure 1a, the mouse fecal microbiota mainly consisted of three classes, *Bacteroidales*, *Lactobacillales*, and *Clostridiales,* regardless of feeding conditions and sampling time. *Lachnospiraceae* and *Streptococcaceae* were dominant (25-30% and 17-45%, respectively) in the mice feces collected before fiber feeding and from the cellulose-fed mice, while in the feces from the inulin-fed mice, the abundance of *Bacteroidaceae* increased relative to other bacteria (17% in morning and 41% in evening), while that of *Lachnospiraceae* decreased (20% in morning and 12% in evening). These relative abundance profiles indicate a microbial composition increment of *Bacteroidaceae* in the feces of inulin-fed mice beyond typical diurnal oscillations in comparison to that in the cellulose-fed mice (p-values <0.005, Tukey’s HSD test).

**Figure 1.**
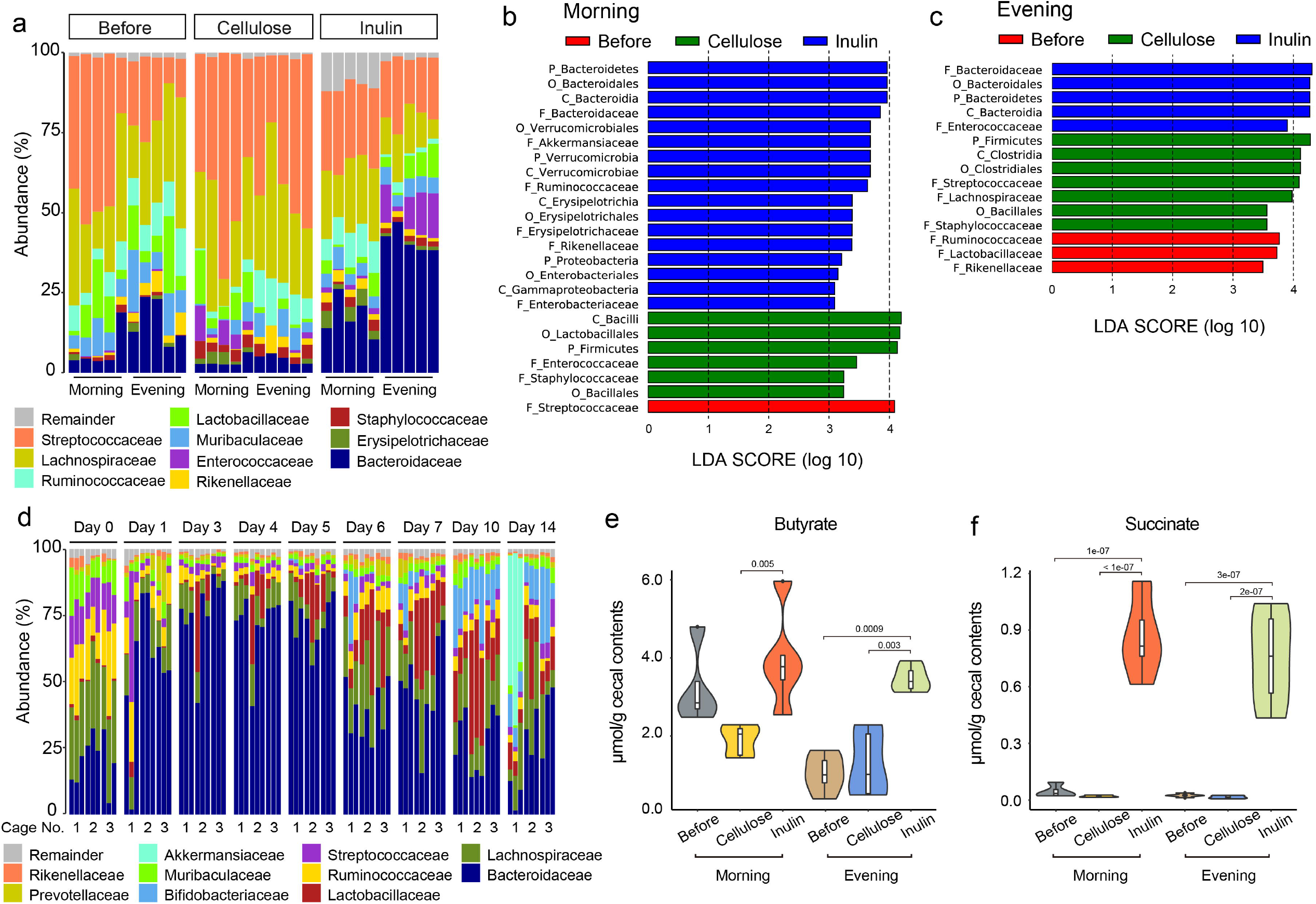
Fecal microbial composition and cecum metabolites in inulin-fed mice. **a.** Relative family-level abundance profiles of mouse fecal microbiomes in the morning and evening before and after two weeks of inulin or cellulose feeding. Box plots indicate the phylogenetic composition of the fecal microbiota samples obtained from each mouse by 16S rRNA gene sequencing (n = 5). **b, c.** Linear discriminant analysis (LDA) scores computed for differentially abundant taxa in the fecal microbiomes of mice before fiber feeding, cellulose-, and inulin-fed mice in the morning (**b**) or evening (**c**) obtained from each mouse by 16S rRNA gene sequencing. The length indicates the effect size associated with each taxon. *P* <0.05 by Wilcoxon signed-rank test; LDA score >2. **d.** Time-dependent changes in relative family-level abundance profiles of the microbiota in each mouse by 16S rRNA gene sequencing. Nine mice in three different cages from a different animal lot from (**a**) were fed inulin for two weeks, and fecal samples were collected in the morning for microbiome composition analysis. Box plots indicate the phylogenetic compositions of fecal microbiota samples from each mouse (n = 3, one from each cage). **e, f.** Concentrations of butyrate (**e**) and succinate (**f**) in the cecum of mice before and after two weeks of inulin or cellulose feeding in the morning or evening. The mice are the same as those shown in (**a**). Violin plots indicate the SCFA concentration in each mouse (n = 5, Tukey’s HSD test).

To identify the specific bacterial taxa associated with inulin fermentation, we compared the fecal microbiota of mice before fiber feeding and inulin-fed mice using the LDA effect size (LEfSe) analysis [20]. Differentially abundant taxa in the fecal microbiota and their predominant bacteria are shown in Figures 1b and 1c. These results primarily indicated the diurnal differences, but revealed that *Bacteroidales* were abundant (LDA score >2, p-value <0.05) in inulin-fed mice at both time points.

To confirm this effect, we monitored time-dependent microbiota composition changes in inulin-fed mice from another animal lot. The feces of inulin-fed mice were sampled in the morning for two weeks. As shown in Figure 1d, *Bacteroidaceae* rapidly increased on day 1, though there were individual differences in microbial composition. On days 3 to 5, *Bacteroidaceae* were most abundant in all mice, which were housed in three different cages. After this rapid increase, *Bacteroidaceae* gradually decreased but maintained their higher percentage in all mice except those in cage 1. In this lot of mice, we observed different microbiota compositional changes compared to the previous experiment, such as an increase in *Bifldobacteriaceae* in all cages, and in *Akkermansiaceae* in cage 1 during days 6 to 14. Despite differences in microbial composition between cages and animal lots, the results strongly suggest that *Bacteroidaceae* is a pivotal responder in the intestinal microenvironment of inulin-fed mice. In addition, we found two main *Bacteroidaceae spp.* (OTU2 and OTU4) in all mice, regardless of the fiber feeding condition. As shown in Additional file 2: Figure S1, OTU2 were rapidly increased after starting inulin feeding and showed significant compositional increment after 2-weeks of continuous inulin feeding (p-values < 4e-02, Tukey’s HSD test). In contrast, OTU4 showed a constant composition during inulin feeding, while they showed significant increment compared to that in the cellulose-fed mice in the evening.

We also evaluated cecum SCFAs to observe the effects of inulin on gut microbiome metabolic function. Figures 1e and 1f show the concentrations of butyrate and succinate in the cecum contents of mice used for 16S rRNA gene sequencing, measured by gas chromatography (GC). Although the total SCFA concentration depended on the diet and sampling time points in mice before fiber feeding and in the cellulose-fed mice (data not shown), samples from inulin-fed mice consistently contained higher butyrate and succinate concentrations.

Taken together, microbial composition and metabolome analyses suggest that inulin feeding changes the microbiome composition to increase *Bacteroides* spp. and microbial metabolic functions, resulting in increased succinate and butyrate concentrations in the mouse intestine.

### Single-cell resolution genome analysis of the fecal microbiome of inulin-fed mice

Using the single-cell sequencing based on the SAG-gel platform (Figure 2a), mouse intestinal microbes were captured in agarose gel beads by microfluidic droplet generator. Cell lysis with an enzyme cocktail, genomic DNA purification [21], and subsequent wholegenome amplification [22, 23] were all performed in the agarose gel matrix. After sorting the single beads, which contain single-cell amplified DNA, into standard PCR plates using a cell sorter, the DNA samples were re-amplified as a single-cell amplified genome (SAG) library. From 464 single-cell captured beads derived from mice fecal microbiota samples before (180 SAGs) and after inulin or cellulose feeding (166 SAGs), a total of 346 SAGs, named IMSAG_001 to 346, were sequenced from the SAG library (Figure 2b). According to the standards developed by the Genomic Standards Consortium [24], we recovered one high-quality draft genome, 43 medium-quality draft genomes, 258 low-quality draft genomes, 38 short genome fragments (< 100 kb or 0% completeness), and six contaminated genomes (Figure 2b and Additional file 1: Table S1). The average completeness and contamination of the draft genomes were 31.8% and 0.6%, respectively, with an average of 296 contigs (Figure 2c). Notably, only 15 of the 346 (4.3%) SAGs had >5% contamination. Approximately 15% of SAGs were classified as high-or medium-quality draft genomes, ranging from 1.3-3.6 Mb in total contig length. The high- and medium-quality draft genomes had an average N50 of 16.4 kb, while the low-quality draft genomes had an average of 10.4 kb (Figure 2d). An average of 17 tRNAs were detected in the high- and medium-quality draft genomes (Figure 2e). Of the 302 SAG draft genomes, 267 (88%) recovered the 16S rRNA gene, 71% were >1,400 bp in size (Figure 2f), and 25% contained an rRNA operon that encoded the 16S, 23S, and 5S rRNA genes in single contigs (Additional file 1: Table S1). When we conducted nucleotide BLAST searches against a database of 16S rRNA gene sequences, 284 SAGs were similar to intestinal microbe 16S rRNA gene sequences with >80% identity, whereas 216 SAGs showed <97% identity with the top search results, suggesting that many SAGs were obtained from uncultured bacteria that have not yet been registered in the database (Additional file 1: Table S2).

**Figure 2.**
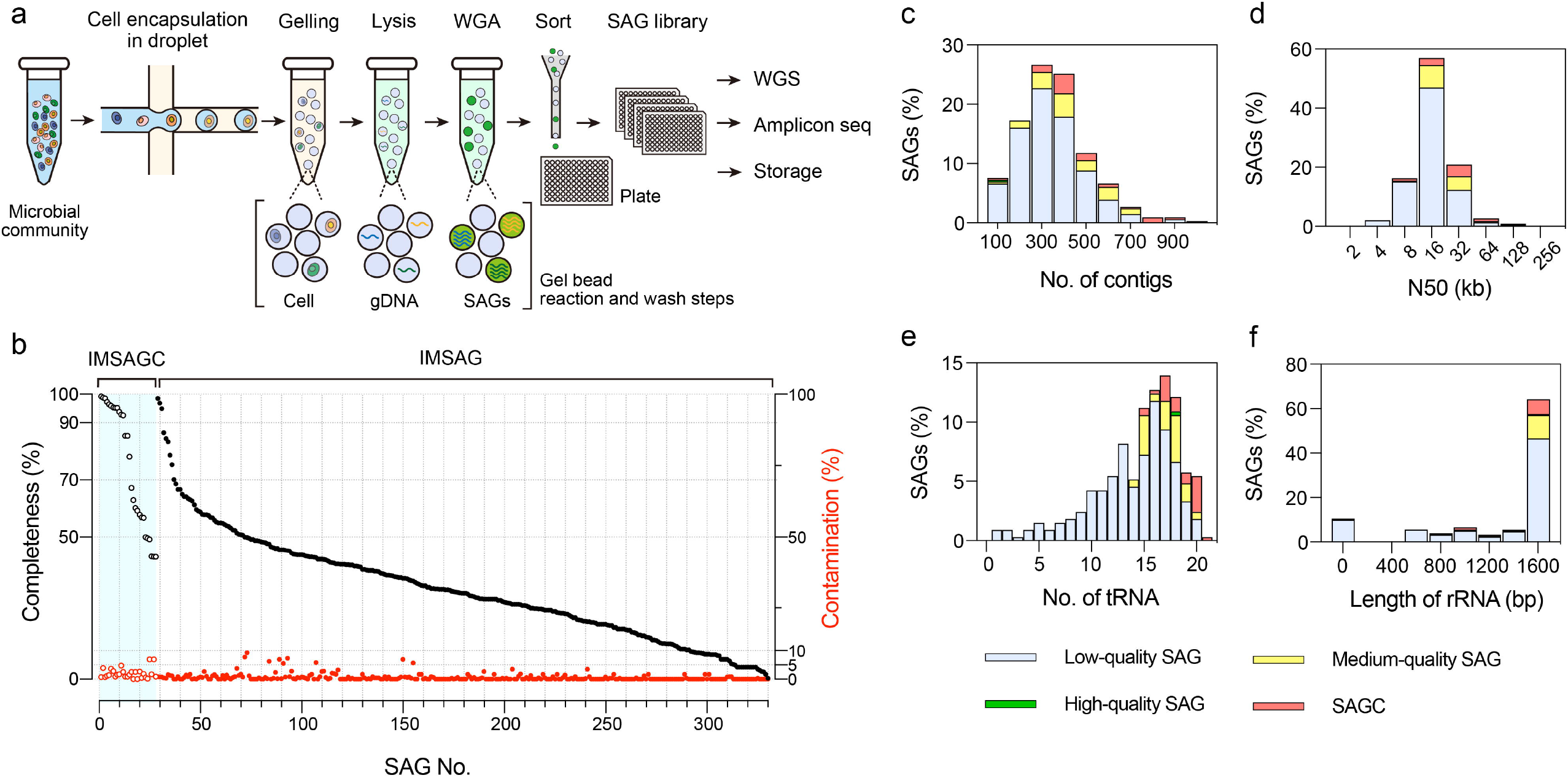
Single-cell genome sequencing of fecal microbiomes in inulin-fed mice. **a.** Workflow for SAG-gel-based single-cell genome sequencing of bacterial cells in a complex microbial community. Individual bacterial cells are randomly captured in picolitersized gel beads and processed by in-gel lysis and whole-genome amplification (WGA). Single-cell amplified genomes (SAGs) in the gel are fluorescently detected and sorted into well plates as a SAG library. The SAGs are re-amplified for further analysis by NGS, amplicon sequencing, and storage. **b-f** Assembly qualities of 324 SAGs, excluding short fragments (length < 100 kb) and contaminated samples (contamination > 10%). **b.** Completeness and contamination statistics for 302 SAGs (IMSAGs) and 22 composite SAGs (IMSAGCs). **c-f.** The number of contigs, N50 values, number of tRNAs, and 16S rRNA gene lengths for 324 SAGs.

Qualified SAGs were classified as 267 bacteria, spanning two phyla, four classes, seven orders, and 14 families with GTDB-Tk [25]. Of all samples, 123 (46%) were classified as *Lachnospiraceae,* followed by *Bacteroidaceae* (65 (24%)), and *Muribaculaceae* (27 (10%); Additional file 1: Table S2). The phylogenetic distribution of these SAGs was slightly different from that of 16S rDNA genes acquired from a metagenomic sample, and the SAGs covered 62 of 298 operational taxonomic units (OTUs; 21%; Additional file 1: Table S3). The SAGs covered seven major bacterial families in the metagenomic analysis (>1% of the total OTUs in the inulin and cellulose samples) but missed the genomes of some grampositive bacteria, such as *Streptococcaceae, Enterococcaceae,* and *Staphylococcaceae*. Moreover, some SAGs had strain marker gene identities ≥99%, and average nucleotide identities (ANIs) ≥95% were co-assembled to improve their completeness as single strain genomes, which were named IMSAGC_001 to 028 (Additional file 1: Table S4). Finally, we obtained 31 different SAG strains (22 composite and 9 raw) that were graded as high-(7) or medium-quality (24) draft genomes among seven families (Table 1 and Additional file 1: Table S5). These included two *Bacteroidaceae* spp. (IMSAGC_001 and IMSAGC_004) that had identical 16S rRNA genes to *Bacteroidaceae* (OTU2 and OTU4, respectively), which were potential inulin-responders in inulin-fed mice feces (Additional file 1: Tables S3 and S5, and Additional file 2: Figure S1).

**Table 1.** Statistics for high and medium quality draft genomes

| SAG ID | Lineage<sup>a</sup> | # contigs | Total length (Mb) | Largest contig (bp) | N50 (bp) | GC (%) | # CDS | Completeness (%)<sup>b</sup> | Contamination (%)<sup>b</sup> | SAG quality<sup>c</sup> |
| --- | --- | --- | --- | --- | --- | --- | --- | --- | --- | --- |
| IMSAGC_001 | Bacteroidota;Bacteroidia;Bacteroidales;Bacteroidaceae;Bacteroides;Bacteroides acidifaciens | 369 | 4.93 | 125378 | 29923 | 43.1 | 4165 | 97.27 | 1.12 | High |
| IMSAGC_002 | Firmicutes_A;Clostridia;Lachnospirales;Lachnospiraceae;28-4 | 709 | 5.08 | 70754 | 13000 | 44.6 | 4725 | 93.87 | 1.78 | Medium |
| IMSAGC_003 | Firmicutes_A;Clostridia;Lachnospirales;Lachnospiraceae;CAG-65 | 308 | 4.62 | 140873 | 48295 | 50.7 | 4203 | 98.85 | 3.97 | Medium |
| IMSAGC_004 | Bacteroidota;Bacteroidia;Bacteroidales;Bacteroidaceae;Bacteroides_B | 305 | 4.22 | 106819 | 34968 | 43.1 | 3598 | 98.62 | 0.8 | Medium |
| IMSAGC_005 | Firmicutes_A;Clostridia;Lachnospirales;Lachnospiraceae;CAG-95 | 317 | 4.23 | 113574 | 39279 | 43.6 | 4110 | 96.45 | 1.45 | Medium |
| IMSAGC_006 | Bacteroidota;Bacteroidia;Bacteroidales;Muribaculaceae;UBA3263;GCA_001689615.1 | 265 | 2.42 | 113083 | 24935 | 50.4 | 2284 | 85.47 | 0.89 | Medium |
| IMSAGC_007 | Firmicutes_A;Clostridia;Lachnospirales;Lachnospiraceae | 496 | 5.06 | 122936 | 25158 | 47.1 | 4957 | 95.89 | 3.59 | Medium |
| IMSAGC_008 | Bacteroidota;Bacteroidia;Bacteroidales;Muribaculaceae;CAG-873 | 380 | 2.66 | 59381 | 16733 | 52.1 | 2436 | 78.2 | 1.58 | Medium |
| IMSAGC_009 | Firmicutes_A;Clostridia;Lachnospirales;Lachnospiraceae;CAG-95 | 597 | 5.02 | 113931 | 23340 | 44.4 | 4633 | 95.33 | 2.87 | Medium |
| IMSAGC_010 | Firmicutes;Bacilli;Lactobacillales;Lactobacillaceae;Lactobacillus;Lactobacillus johnsonii | 54 | 1.91 | 214598 | 104736 | 34.3 | 1789 | 99.22 | 0.78 | High |
| IMSAGC_011 | Firmicutes_A;Clostridia;Lachnospirales;Lachnospiraceae;COE1 | 431 | 4.25 | 143151 | 27757 | 37.9 | 3664 | 92.66 | 2.59 | High |
| IMSAGC_012 | Firmicutes_A;Clostridia;Lachnospirales;Lachnospiraceae;Eubacterium_J | 419 | 3.92 | 117373 | 25561 | 45.0 | 3748 | 92.92 | 4.75 | Medium |
| IMSAGC_013 | Firmicutes_A;Clostridia;Lachnospirales;Lachnospiraceae;14-2 | 884 | 5.14 | 67964 | 11980 | 44.6 | 4945 | 85.42 | 0.95 | Medium |
| IMSAGC_014 | Bacteroidota;Bacteroidia;Bacteroidales;Bacteroidaceae;F0040 | 273 | 2.64 | 87270 | 20006 | 47.1 | 2340 | 95.18 | 1.58 | Medium |
| IMSAGC_015 | Firmicutes_A;Clostridia;Lachnospirales;Lachnospiraceae;Dorea | 388 | 2.55 | 63422 | 12891 | 45.3 | 2429 | 67.29 | 2.53 | Medium |
| IMSAGC_016 | Bacteroidota;Bacteroidia;Bacteroidales;Muribaculaceae;CAG-485 | 385 | 1.94 | 50916 | 8558 | 46.4 | 1735 | 56.91 | 0.5 | Medium |
| IMSAGC_017 | Firmicutes;Bacilli;Erysipelotrichales;Erysipelatoclostridiaceae;Erysipelatoclostridium;Erysipelatoclostridium cocleatum | 299 | 2.69 | 71470 | 19801 | 28.9 | 2384 | 95.28 | 0.94 | Medium |
| IMSAGC_019 | Firmicutes_A;Clostridia;Lachnospirales;Lachnospiraceae;Dorea | 311 | 2.42 | 198629 | 20896 | 44.1 | 2370 | 62.98 | 0 | Medium |
| IMSAGC_020 | Firmicutes_A;Clostridia;Lachnospirales;Lachnospiraceae;CAG-95 | 756 | 4.13 | 136678 | 11653 | 45.8 | 4150 | 57.89 | 2.63 | Medium |
| IMSAGC_021 | Bacteria;Firmicutes_A;Clostridia;Lachnospirales;Lachnospiraceae;Dorea | 642 | 3.03 | 69412 | 9299 | 45.8 | 2954 | 60.15 | 2.58 | Medium |
| IMSAGC_027 | Bacteroidota;Bacteroidia;Bacteroidales;Muribaculaceae;CAG-1031;GCA_001689585.1 | 314 | 2.00 | 63820 | 15566 | 50.5 | 1738 | 59.25 | 0 | Medium |
| IMSAGC_028 | Bacteroidota;Bacteroidia;Bacteroidales;Rikenellaceae;Alistipes | 265 | 1.73 | 77838 | 22105 | 52.5 | 1581 | 56.76 | 1.92 | Medium |
| IMSAG_013 | Firmicutes_A;Clostridia;Oscillospirales;CAG-272 | 229 | 1.72 | 93908 | 16943 | 45.4 | 1622 | 59.65 | 0 | Medium |
| IMSAG_025 | Bacteroidota;Bacteroidia;Bacteroidales;Muribaculaceae;CAG-485 | 667 | 2.51 | 52448 | 6077 | 45.3 | 2558 | 50 | 7.76 | Medium |
| IMSAG_044 | Firmicutes;Bacilli;Lactobacillales;Lactobacillaceae;Lactobacillus_H;Lactobacillus_H reuteri_A | 200 | 1.82 | 108153 | 17958 | 38.3 | 1735 | 86.54 | 0.54 | Medium |
| IMSAG_049 | Firmicutes_A;Clostridia;Lachnospirales | 317 | 1.74 | 50322 | 11294 | 39.6 | 1762 | 54.88 | 1.34 | Medium |
| IMSAG_117 | Firmicutes;Bacilli;Lactobacillales;Lactobacillaceae;Lactobacillus_B;Lactobacillus_B murinus | 314 | 1.86 | 38237 | 10806 | 40.5 | 1824 | 78.75 | 1.31 | Medium |
| IMSAG_185 | Firmicutes_A;Clostridia;Lachnospirales;Lachnospiraceae;CAG-65 | 317 | 2.25 | 78895 | 17758 | 49.9 | 2117 | 54.01 | 1.45 | Medium |
| IMSAG_192 | Bacteroidota;Bacteroidia;Bacteroidales;Muribaculaceae;Muribaculum;Muribaculum intestinale | 342 | 1.89 | 59599 | 11500 | 49.3 | 1711 | 57.55 | 0 | Medium |
| IMSAG_249 | Firmicutes_A;Clostridia;Lachnospirales;Lachnospiraceae;CAG-95 | 428 | 2.83 | 108242 | 17278 | 44.4 | 2659 | 55.18 | 0.91 | Medium |
| IMSAG_250 | Firmicutes_A;Clostridia;Oscillospirales;DTU089;Eubacterium_R | 147 | 1.35 | 87509 | 28237 | 37.8 | 1309 | 68.68 | 0 | Medium |

<sup>a</sup> Assigned using GTDB-Tk.
<sup>b</sup> Estimated using CheckM.
<sup>c</sup> Classified according to GSC MISAG.

### Genomic features of *Bacteroides* spp. draft genomes

*Bacteroides* spp. are capable of metabolizing fructans, a class of plant and microbial polysaccharides [7, 9, 11]. The completeness of the inulin responder *Bacteroides* draft genomes IMSAGC_001 and IMSAGC_004 were sufficiently high (≥97%) for comparison with publicly available genomes of *Bacteroides* spp. known to be fructan utilizers or showing similarity to these genomes (Additional file 1: Table S6 and Figure 3a). Both had >4.2 Mb genome sizes, 43% GC content, and >3,500 coding sequences (CDSs), similar to the genomes of *Bacteroides* spp. including *B. thetaiotaomicron* (Bt), *B. ovatus* (Bo), *B. caccae* (Bc), *B. vulgatus* (Bv), *B. uniformis* (Bu), *B. fragilis* (Bf), and *B. acidifaciens* (Ba; Figure 3a and Table S6). Genes in these *Bacteroides* groups were clustered into 5,013 different orthogroups. The orthogroup distributions between different *Bacteroides* group members are shown in Figure 3b; of these orthogroups, 1,150 were defined as core orthogroups. IMSAGC_001 and IMSAGC_004 shared a number of different orthogroups with specific strains, such as Ba (196 orthogroups) and Bv (99 orthogroups), consistent with the phylogenetic analysis (Figure 3a). Although the vast majority of the orthogroups (151 and 57 in IMSAGC_001 and IMSAGC_004, respectively) were classified as hypothetical proteins of unknown function, most orthogroups shared between IMSAGC_001 and Ba belonged to category O (carbohydrate transport and metabolism; nine orthogroups), followed by categories T and P (signal transduction mechanisms and inorganic ion transport and metabolism, respectively; six orthogroups each). In addition, IMSAGC_001 and IMSAGC 004 shared 47 unique orthogroups (37 with unknown functions).

**Figure 3.**
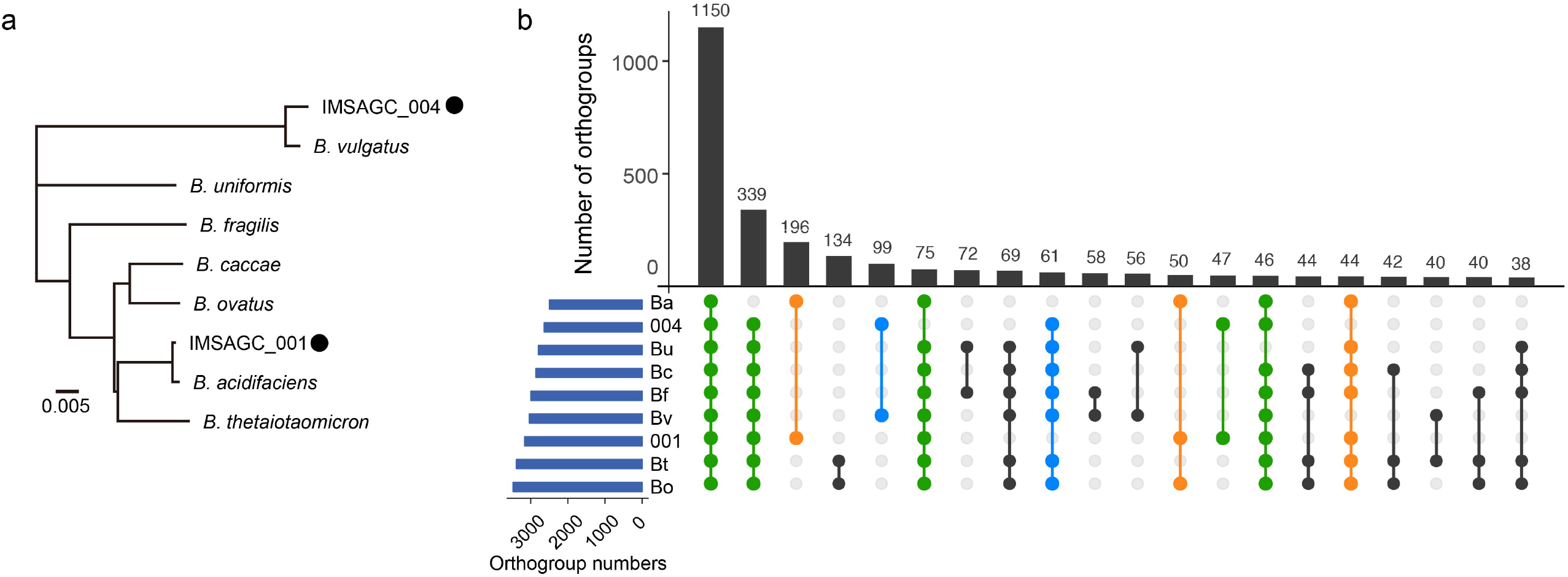
Comparative analysis of the inulin responder *Bacteroides* genomes IMSAGC_001 and IMSAGC_004. **a**. Phylogenetic tree of inulin-associated *Bacteroides* spp. found in this study and reference *Bacteroides* strains. The tree is based on the alignment of concatenated amino acid sequences of collocated sets of ubiquitous, single-copy genes within a phylogenetic lineage, as used in CheckM analysis. IMSAGC_001 and IMSAGC_004 are indicated by black circles. **b**. UpSet plot comparing shared orthogroup counts between reference *Bacteroides* strains. Orthogroups shared between IMSAGC_001 and other *Bacteroides* group members are colored in orange, whereas orthogroups shared between IMSAGC_004 and other *Bacteroides* group members are colored in blue. Orthogroups shared between IMSAGC**_**001, IMSAGC_004, and other members are colored in green. Bt, *B. thetaiotaomicron*; Bo, *B. ovatus;* Bc, *B. caccae*; Bv, *B. vulgatus*; Bu, *B. uniformis*; Bf, *B. fragilis*; Ba, *B. acidifaciens.*

### Identification of putative polysaccharide utilization loci from *Bacteroides* spp. draft genomes

Most of the glycan degrading and import machinery within *Bacteroides* genomes are encoded within clusters of coregulated genes known as polysaccharide utilization loci (PUL). The defining characteristic of a PUL is the presence of a pair of genes homologous to *susD* and *susC*, which encode outer membrane proteins that bind inulin and import its digestion products [8, 11, 26]. The *susC* and *susD* homologs are usually associated with genes encoding the machinery necessary to convert extracellular polysaccharides into intracellular monosaccharides, such as glycoside hydrolases (GHs) or polysaccharide lyases (PLs). In addition, most PULs contain or are closely linked to genes encoding an inner membrane-associated sensor-regulator system, including the hybrid two-component system. Notably, some *Bacteroides* species, such as Bc, Bo, Bt, and Bu, can utilize inulin with similar efficiency to glucose [7, 8, 11], while Bv lacks *susC-* and *susD*-like genes within its fructan PUL and cannot use inulin. Bt and Bo encode a number of predicted signal peptidase (SP)II-containing GHs/PLs and act as public goods producers in various networks of polysaccharide utilizers [7]. These producer-derived GHs are extracellularly transported to create polysaccharide degradation products that can be utilized for growth of concomitant recipients, such as Bv, at considerable distance from the producer.

We detected PULs containing inulinases in two *Bacteroides* draft genomes (IMSAGC_001 and IMSAGC_004; Figures 4a and 4b). We also identified other *Bacteroidales*, such as IMSAGC_014 and IMSAGC_006, that had no PULs encoding inulinases. At the whole genome level, IMSAGC_001 showed similarity to inulin utilizers such as Bo, Bc, and Bt, while IMSAGC_004 was similar to the inulin non-utilizer Bv (Figure 3a). In IMSAGC_001, the sequences of GH genes, including inulinase, showed high similarity to those of Bc and Bo, which have superior ability to use inulin [7, 11] and had Q highest similarity to Ba (Figure 4c). IMSAGC_004 also contained a GH gene similar to that of known inulin utilizers. The GHs of IMSAGC_001 and IMSAGC_004 are predicted to be periplasmic, as they contain SPI cleavage sites. In addition, IMSAGC_001, IMSAGC_004, and Ba have no predicted SPII-containing GHs and PLs in their inulin utilization loci (Figure 4b). Since *Bacteroides* spp. show differences in numbers of GH/PL-encoding genes and uniformity among strains of a given species [7], we compared the number of SPII-containing GH/PLs among *Bacteroides* genomes. IMSAGC_001 had a medium level of SPII-containing GH/PLs (56, Additional file 1: Table S7), while IMSAGC_004 and Ba had extremely low levels of SPII-containing GH/PLs (26 and 32, respectively). This result indicated that IMSAGC_001 and IMSAGC_004 might contain different types of machinery for polysaccharide utilization from Bt and Bo, which have several SPII-containing GH/PLs, which act as producers for a larger repertoire of polysaccharides.

**Figure 4.**
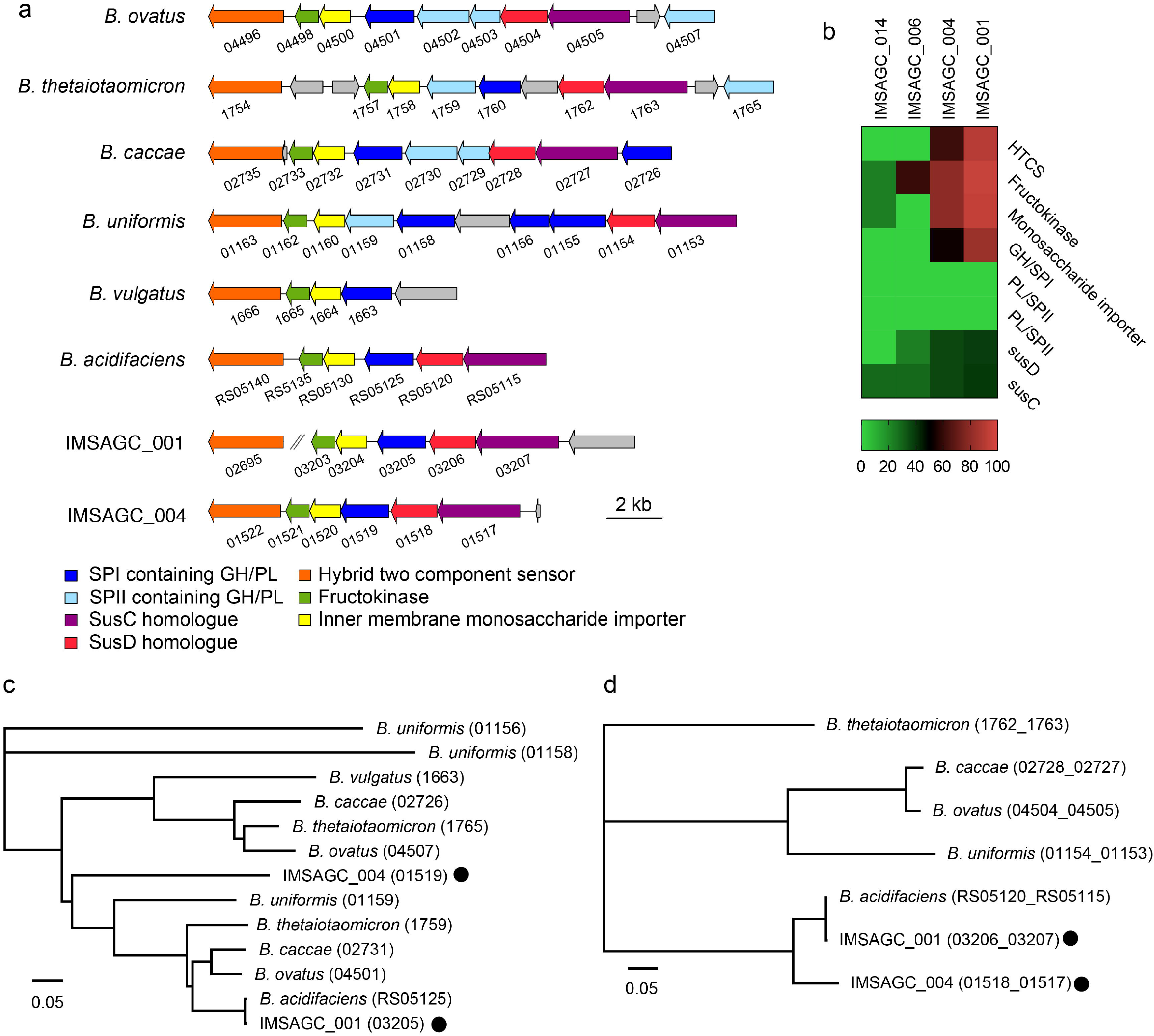
Identification of putative inulin utilization loci in the inulin responder *Bacteroides* genomes IMSAGC_001 and IMSAGC_004. **a.** Polysaccharide utilization loci (PUL) from reference *Bacteroides* species, IMSAGC**_**001, and IMSAGC**_**004. Common predicted functions are color-coded while intervening unrelated genes are in gray. Signal peptide, SP; glycoside hydrolase, GH; polysaccharide lyase, PL. **b.** Heat map of the identities of amino acid sequences in PUL genes between inulin responder (001 and 004) and non-responder (006 and 014) *Bacteroidales* and the known inulin utilizer *B. ovatus.* **c, d.** Phylogenetic analysis of glycoside hydrolase (**c**) and a concatenated sequence containing *SusC* and *SusD* (**d**) between reference *Bacteroides* species, IMSAGC**_**001, and IMSAGC**_**004. IMSAGC**_**001 and IMSAGC**_**004 are indicated with black circles.

For *susC/D* homologs, IMSAGC_001 and IMSAGC_004 clustered with Ba and apart from other inulin utilizers (Bc, Bo, Bu, and Bt; Figure 4d). Homologs of the SusC and SusD outer membrane proteins are the defining features of the Sus-system, and genes encoding these proteins are found as a pair in most *Bacteroidetes* genomes. Inulin chains that may or may not be extracellularly cleaved are threaded through the SusC porin, and all degradation occurs in the periplasm. In Bo, deletion of the *susC* and *susD* orthologs results in significant growth impairment on inulin, and the orthologs are predicted to function in inulin binding and import without prior extracellular digestion [9, 26]. As our data suggested that IMSAGC_001 and IMSAGC_004 had low similarity to the Bo *susC/D* homologs, we next compared genome sequence similarities through structure prediction-based analysis. Based on structure prediction, the susC/D homologs in IMSAGC_001 and IMSAGC_004 could form complex structures composed of homodimers of an extracellular SusD-like lipoprotein and an integral membrane SusC-like transporter (Additional file 2: Figure S2). These predicted structures showed high similarity to the Ba complex at the amino acid level (Additional file 2: Figure S2).

In summary, *Bacteroides* IMSAGC_001, which were significantly increased in inulin-fed mouse feces, and IMSAGC_04, which showed no apparent changes, possess putative PULs that include *susC/D* homolog pairs, a putative outer membrane lipoprotein, a putative inner membrane monosaccharide importer, a putative fructokinase, and a putative inulinspecific GH. No other SAGs had higher similarity to *Bacteroides* PUL. We predict that these bacteria import inulin directly via SusC and SusD for degradation to monosaccharides in the periplasm without prior extracellular degradation.

### Identification of metabolic characters in inulin-utilizing strains and other bacterial SAGs

Having identified key responder bacterial strains for inulin metabolism, we next examined their fundamental metabolic functions in the mouse gut microbiome. Additional file 1: Tables S8 and S9 summarize the metabolic functions of all high- and medium-quality genomes obtained in this study (IMSAGC series with >80% completeness). Notably, inulin-fed mice displayed significant increases in *Bacteroides* in their microbial composition and succinate and butyrate concentrations in their SCFAs. For succinate, we focused on the reductive tricarboxylic acid (TCA) cycle, as it produces the major end product of energy metabolism in anaerobic microorganisms. Many bacteria have incomplete TCA cycles and are commensal obligate anaerobes, including *Bacteroides* spp., which use a branched TCA cycle to support fumarate respiration [27, 28]. The end product of fumarate reduction, succinate, is secreted into the extracellular environment. In IMSAGC_001 and IMSAGC_004, we detected reductive and incomplete TCA cycles from oxaloacetate to succinate (Additional file 2: Figure S3 and Additional file 1: Table S8), while no other predominant SAG contained both an inulin PUL and a succinate pathway.

In single-cell genome sequencing, the profiling of SAG metabolic pathways provides various insights into the traits of diverse microbiota at once, because each metabolic gene is specifically assigned to specific bacterial strains (Additional file 1: Table S9). As an interesting functional signature, we detected the presence of antibiotic resistance modules, such as vancomycin resistance, D-Ala-D-Ser type, in specific *Lachnospiraceae* strains (IMSAGC_003 and IMSAGC_007). We observed differences in conserved biosynthetic pathways in cofactor and vitamin metabolism, such as biotin and cobalamin synthesis, among different intestinal microbes. Comparing IMSAGC_001, IMSAGC_004, and other *Bacteroides* spp., IMSAGC_004 lacks proline and tryptophan biosynthesis pathways, as well as the first carbon oxidation pathway in the TCA cycle, but has a conserved cobalamin biosynthetic pathway similar to those of Bf, Bu, and Bv. Both IMSAGC_001 and IMSAGC_004 lack glycine betaine/proline transport systems, similar to Ba and Bu. The inulin nonresponsive *Bacteroidales* spp. (IMSAGC_014 and IMSAGC_006) lacked some carbohydrate metabolism pathways and saccharide transport systems compared to known polysaccharide utilizers. These results suggest that inulin-responsive *Bacteroides* IMSAGC_001, IMSAGC_004, and inulin non-responder *Bacteroidales* spp. play different metabolic roles in the mouse gut microbiota.

## Discussion

Inulin (β2-1 fructan) is resistant to host-mediated digestion, resulting in remarkable fecal microbiome changes and cecum SCFA concentrations through intestinal microbial fermentation. In this study, to elucidate the inulin-responders in mouse intestine microbiota, we conducted 16S rRNA gene sequencing for the assessment of changes in the inulindependent microbiota composition, metabolome analysis for the evaluation of SCFA production, and single-cell genome sequencing for obtaining the draft genomes of inulin-responders. The microbiome composition and SCFA amounts displayed typical diurnal oscillations, and inulin intake resulted in significant changes beyond these. *Bacteroides* spp. are known to utilize levan and inulin (Figures 1 and 3, Additional file 1: Table S6). Microbiome composition monitoring demonstrated rapid responses by *Bacteroides* IMSAGC_001 to inulin supplementation and dynamic daily changes. However, reference *Bacteroides* strains have different origins and uncultured strains present in individual hosts could have different metabolic features. *Bacteroides* members have differing abilities to use these various plant polysaccharides. Species that utilize a particular polysaccharide (producers), such as Bo, liberate polysaccharide degradation products that are consumed by other recipients unable to grow using the intact polysaccharide [7, 8].

Functional characterization of bacteria using 16S rRNA gene sequencing databases may lead to misinterpretations. From a partial 16S rRNA gene sequence, responder bacteria can only be identified at the family level, *Bacteroidaceae*, and insights on the inulin utilization abilities of specific bacteria in the diverse microbial community cannot be obtained (Figure 1 and Additional file 1: Table S3). Therefore, single-cell sequencing was used to analyze closely related species and compare uncultured bacterial genomes to identify inulin-responders in complex microbial communities.

The advantage of single-cell sequencing with the SAG-gel platform is that it can select specific sequencing samples after generating large numbers of SAGs. Since we can adjust the total sequencing effort according to the number of SAGs required, it is not necessary to produce a large number of reads to cover target bacteria with the same depth as in shotgun metagenomic sequencing. Since each SAG is composed of tens of thousands of reads on average, limiting the analysis to specific strains enables extremely cost-efficient whole-genome sequencing of target cells. Indeed, approximately half of the SAGs were obtained from up to 96-plex indexes in single sequence runs by MiSeq (total ~3Gb). This prescreening can utilize marker genes, such as 16S rRNA genes, to select *Bacteroidaceae* and other abundant strains based on initial 16S rRNA gene sequencing. Even if the target bacteria were selected from the results of initial 16S rRNA gene sequencing, it is challenging to demonstrate direct agreement between the MAGs and the target bacteria as most MAGs lack the rRNA and tRNA sequences in the contigs [15–17]. Thus, SAGs with functional and phylogenetic information are useful in combination with stepwise multilayer analyses, including screening through 16S rRNA gene sequencing, to achieve the strategic genomic analysis of target bacteria.

Conversely, nearly 90% of the SAGs obtained in this study contained 16S rRNA gene sequences, 70% of which were full-length, and many existed in operons with the 5S and 23S rRNA genes. The SAGs provide the genetic information of dominant microbes by covering dominant OTUs existing in >1% of the fecal microbiome community. However, there is still room for improvement. For example, modifying the cell lysis procedure could enhance the variety of SAGs detected, including gram-positive bacteria. In 16S rRNA gene homology analysis, a considerable number of SAGs showed low similarity to reference genomes, suggesting that these SAGs contain the genomes of species currently undescribed in public databases. Furthermore, by acquiring duplicate SAG data from identical strains, we could assemble the draft genomes using a cleaning and co-assembly strategy [22, 23]. For example, IMSAGC_001 and IMSAGC_004 were duplicated 41 and 12 times, respectively, allowing the construction of composite draft genomes, which are very advantageous for obtaining genomes of medium or high quality. These two main Bacteroides *spp.* were present in all mice, regardless of inulin feeding (Table S3). Thus, we utilized raw SAGs for assembly improvement from all mice regardless of fiber feeding conditions. According to the definition used in Almeida et al. [16], 40% of the draft genomes of quality higher than medium were qualified as near-complete (>90% completeness and < 5% contamination, Additional file 1:Table S5). Hence, the SAG-gel platform can provide increased numbers of draft genomes from single microbial cells, with higher completeness than other single-cell genome sequencing techniques [22, 23, 29–32]. The ratio of high- and medium-quality draft genomes was comparable with recent metagenomic assembly and binning strategies for obtaining MAGs as draft genomes [15–17].

Pan-genome analysis within the same environment reveals what bacteria are present and which harbor specific genes (Additional file 1:Table S9). Our results revealed PUL and succinate synthesis pathways in inulin responder *Bacteroidaceae*. In addition, comparative SAG analysis allowed us to understand the conservation of various pathways, such as those involved in antibiotic resistance and vitamin synthesis, in each bacterial strain in the intestinal microbiome. Single strain-resolution fingerprinting of the microbiome enables the identification of bacteria or genes responsible for specific metabolic function from a complex microbial community. In this study, two responder *Bacteroides* genomes had inulin-utilizing PUL clusters that showed similarity to known inulin utilizers. Our genome analyses suggested that *Bacteroides* IMSAGC_001 and IMSAGC_004, which can take up and degrade inulin, grow predominantly in the intestines of inulin-fed mice, and produce succinate through the glycolytic system and a reductive partial TCA cycle for release into the intestinal environment. We have identified the potential inulin utilizers *Bacteroides* IMSAGC_001 and IMSAGC_004 in uncultured fecal microbiomes as a new subgroup with Ba. Ba was originally isolated from mouse cecum as two strains that differed in their starch hydrolysis functions [33]. Their major end products of glucose fermentation are acetate and succinate, similar to those predicted for *Bacteroides* IMSAGC_001 and IMSAGC_004. Our data suggest that *Bacteroides* IMSAGC_001 and IMSAGC_004 utilize inulin for their metabolism and lack the ability to liberate the fructose for other species in the same manner as Bf and Ba [7]. The significant increase in *Bacteroides* IMSAGC_001 over IMSAGC_004 (Additional file: Figure S1) may be due to differences in the cell growth rate, as well as inulin uptake and degradation rates, given the differences in GHs and other PUL genes.

## Conclusion

In this study, single-cell genomics enabled detailed microbiome analysis and identified inulin-responders with putative roles in dietary fiber metabolism. The technique generates a massive preparation of SAGs, enabling the functional analysis of uncultured bacteria in the intestinal microbiome and the consideration of host responses. The analysis makes it possible to estimate metabolic lineages in bacterial PUL and metabolic outcomes in the intestinal environment, such as SCFA production, based on ingested fibers. Genetic and functional differences between intestinal bacterial species are predictive of *in vivo* competitiveness in the presence of dietary fibers. Taxonomic and functional genes regulating bacterial metabolic capacity can serve as potential biomarkers in microbiome datasets and can be sequenced to assess microbiota control via diet. In the future, it may be possible to estimate responders in advance based on the presence of these microorganisms and genes.

## Methods

### Mice

Wild-type six-week-old male mice from the BALB/c background (Tokyo Laboratory Animals Science Co., Japan) were used. All mice were housed in a 12:12 light:darkness cycle (Zeitgeber time (ZT) 0 = 8 AM, ZT 12 = 8 PM). Before feeding experiments, the mice were fed AIN93M (Oriental Yeast Co., Japan) for one week. After that, the mice were fed inulin- or cellulose-supplemented diets for two weeks in the morning or evening. To prepare the fiber-supplemented feed, AIN93M powder was mixed with 5% inulin or 5% cellulose, and then pellets were formed with tap water. Fecal samples were collected for 16S rRNA gene sequencing and single-cell sequencing, and cecum samples were harvested before and after the two-week fiber feeding for metabolomic analysis. All samples were stored at −80 °C.

### DNA extraction from fecal samples

Fecal samples (200 mg) were homogenized with 20 mL phosphate-buffered saline (PBS) with a Pellet Pestle™ Cordless Motor (Fisher Scientific). The fecal suspensions were then filtered through 100-μm nylon mesh to remove eukaryotic cells and other debris, and the flow-through suspensions were collected as the bacterial cell fraction. Filtered fecal suspensions were then centrifuged at 9,000 × *g* for 10 min at 4 °C, and the bacterial pellets were resuspended in 800 μL Tris-ethylenediaminetetraacetic acid (TE) 10 buffer (10 mM Tris-HCl/10 mM EDTA). Bacterial pellets were serially incubated with 100 μL lysozyme (150 mg/mL) at 37 °C for 1 h, 20 μL achromopeptidase (100 U/μL) at 37 °C for 30 min, and 50 μL proteinase K (20 mg/mL) with 50 μL 20% sodium dodecyl sulfate (SDS) at 55 °C for 1 h. The lysates were then treated with equal volumes of phenol:chloroform:isoamyl alcohol and centrifuged at 6,000 × *g* for 10 min. After centrifugation, DNA in the upper aqueous layer was transferred to another tube and then pelleted by adding 3M sodium acetate solution and isopropanol, followed by centrifugation at 6,000 × *g* for 10 min. DNA pellets were rinsed with 70% ethanol, dried, and dissolved in TE buffer (10 mM Tris-HCl/1 mM EDTA). The collected DNA samples were purified with 10 μg/mL RNase and incubated at 37 °C for 30 min and precipitated by adding equal volumes of 20% polyethylene glycol solution (PEG6000). After centrifugation at 10,000 × *g* for 10 min at 4 °C. DNA pellets were rinsed with 70% ethanol, dried, and dissolved in TE buffer. The extracted DNA samples were stored at −20 °C.

### 16S rRNA gene sequencing and analysis

The V3–V4 hypervariable regions of 16S rRNA genes were analyzed according to the Illumina protocol for 16S Metagenomic Sequencing Library Preparation. Barcoded amplicons, amplified with 341F and 806R primers (Forward, 5’-TCGTCGGCAGCGTCAGATGTGTATAAGAGACAGCCTACGGGNGGCWGCAG-3’; reverse, 5’-GTCTCGTGGGCTCGGAGATGTGTATAAGAGACAGGACTACHVGGGTATCTAATC C-3’) were sequenced using the Illumina MiSeq 2 × 300 bp platform with MiSeq Reagent Kit v3 (Illumina Co.) according to the manufacturer’s instructions. Raw reads were processed using QIIME2 v.2019.1 [34] using the DADA2 [35] plugin to denoise quality filter reads, call amplicon sequence variants (AS Vs), and generate a feature table of ASV counts and host metadata. In the quality filtering step, the datasets were truncated to read length of 270 to 250 base pairs for the forward and reverse reads (all other parameters were set to default values). After quality filtering, bacterial taxonomies were assigned to the ASV feature table using the Naïve Bayesian Q2 feature classifier as implemented in QIIME2. We compared the data against a SILVA reference database [36] trained on the V3-V4 region of the 16S rRNA gene.

Merged paired-end sequence reads of each sample were trimmed from the primer sequences, and low-quality reads (base error rate >1%) and chimeric sequences were removed using USEARCH [37]. Filtered sequence reads were assigned to operational taxonomic units (OTUs) by open-reference OTU picking at 97% identity using UPARSE algorithm [38] against a SILVA reference database [36]. Taxonomic assignment of the reference sequence was used as the taxonomy for each OTU, and taxonomy table at the family level was generated.

LEfSe analysis was performed to identify taxa displaying the largest differences in microbiota abundance between groups [20]. Only taxa with LDA scores >2.0 and *p*□ <0.05, as determined by Wilcoxon signed-rank test, are shown. All data analyses were performed using R software v3.5.3.

### Metabolome analysis from cecum samples

SCFA concentrations of cecum contents were determined by GC-mass spectrometry. In brief, 50 mg cecum contents were homogenized with ether. After vigorous shaking using a Shake Master Neo (Bio-Medical Science) at 1,500 rpm for 10 min, homogenates were centrifuged at 1,000 × *g* for 10 min, then the top ether layers of each sample were collected and transferred to new glass vials. Aliquots of the ether extracts (80 μL) were mixed with 16 mL N-tert-butyldimethylsilyl-N-methyltrifluoroacetamide. The vials were sealed tightly and heated at 80 °C for 30 min in a water bath. Derivatized samples were loaded onto a 6890N Network GC System (Agilent Technologies) equipped with an HP-5ms column (0.25 mm × 30 m × 0.25 mm) and a 5973 Network Mass Selective Detector (Agilent Technologies). The temperature program used was 60 °C for 3 min, 5 °C/min increases up to 120 °C, then 20 °C/min increases up to 300 °C. Each sample (1.0 mL) was injected with a run time of 30 min. SCFA concentrations were quantified by comparing their peak areas with standards.

### SAG-gel processing step 1: single bacterial cell isolation with gel beads

Microfluidic droplet generators were fabricated and used for in-bead bacterial genome amplification sequencing. After homogenization of mouse feces in PBS, the supernatant was recovered by centrifugation at 2,000 × *g* for 2 s, followed by 15,000 × *g* for 3 min. The resulting cell pellets were suspended in PBS, filtered through 35-μm nylon mesh, washed twice, and resuspended in 1.5% agarose in Dulbecco’s PBS (DPBS; Thermo Fisher Scientific).

Prior to single-cell encapsulation, cell suspensions were adjusted to 0.1 cells/bead to prevent the encapsulation of multiple cells in single beads. Using the droplet generator, single microbial cells were encapsulated in droplets and collected in a 1.5 mL tube, which was chilled on ice for 15 min to form the gel matrix. After solidification, collected droplets were broken with 1H,1H,2H,2H-perfluoro-1-octanol to collect the beads. Then, the gel beads were washed with 500 μL acetone (Sigma), and the solution was mixed vigorously and centrifuged. The acetone supernatant was removed, and 500 μL isopropanol (Sigma) was added, and the solution was mixed vigorously and centrifuged. The isopropanol supernatant was removed, and the gel beads were washed three times with 500 μL DPBS.

### SAG-gel processing step 2: massively parallel single-cell genome amplification

Next, individual cells in beads were lysed by submerging the gel beads in lysis solutions: first, 50 U/μL Ready-lyse Lysozyme Solution (Epicentre), 2 U/mL Zymolyase (Zymo research), 22 U/mL lysostaphin (MERCK), and 250 U/mL mutanolysin (MERCK) in DPBS at 37 °C overnight; second, 0.5 mg/mL achromopeptidase (MERCK) in DPBS at 37 °C for 8 h; and third, 1 mg/mL Proteinase K (Promega) with 0.5% SDS in DPBS at 40 °C overnight. At each reagent replacement step, the gel beads were washed three times with DPBS and then resuspended in the next solution. After lysis, the gel beads were washed with DPBS five times and the supernatant was removed. Then, the beads were suspended in Buffer D2 and subjected to multiple displacement amplification using a REPLI-g Single Cell Kit (QIAGEN).

After WGA at 30 °C for 2 h and 65 °C for 3 min, the gel beads were washed three times with 500 μL DPBS. Then, the beads were stained with 1 × SYBR Green in DPBS. After confirming DNA amplification by the presence of green fluorescence in the gel, fluorescence-positive beads were sorted into 0.8 μL DPBS in 96-well plates by a FACSMelody cell sorter (BD Bioscience) equipped with a 488 nm excitation laser. Green fluorescence intensities were detected through a 550-nm long-pass dichroic filter and a 525-nm band-pass filter using fluorescence detector 1 (FL1) with a PMT voltage of 650 and a logarithmic gain. The sample flow rate was adjusted to approximately 200 events/s. At least 5,000 particles were analyzed for each histogram. After droplet sorting, 96-well plates proceeded through a second round of WGA or were stored at −30 °C.

### SAG-gel processing step 3: multiplex single-cell genome sequencing

After gel bead collection in 96-well plates, second-round MDA was performed with the REPLI-g Single Cell Kit. Buffer D2 (0.6 μL) was added to each well and incubated at 65 °C for 10 min. Then, 8.6 μL of MDA mixture was added and incubated at 30 °C for 120 min. The MDA reaction was terminated by heating at 65 °C for 3 min. After second-round amplification, master library plates of single amplified genomes (SAGs) were prepared. For quality control, aliquots of the SAGs were transferred to replica plates, which were used for DNA yield quantification using a Qubit dsDNA High Sensitivity Assay Kit (Thermo Fisher Scientific) and 16S rRNA gene Sanger sequencing (FASMAC) with the V3-V4 primers described above. Based on the DNA amount and 16S rRNA gene sequence, after removing the contaminated samples, to perform whole-genome sequencing, we screened the SAG samples that had enough DNA for library preparation with qualified 16S rRNA sequences obtained by Sanger sequencing (more than 400 bp and HQ 40% were assessed using the Geneious software (Biomatters, Ltd.)). For sequencing analysis, Illumina libraries were prepared using amplicons from the second-round MDA product, using the Nextera XT DNA Library Prep Kit (Illumina) according to the manufacturer’s instructions. Each SAG library was sequenced using an Illumina MiSeq 2 × 75bp or HiSeq 2 × 150bp configuration (GENEWIZ).

### Single-cell genome sequencing analysis

The sequence reads were assembled *de novo* using SPAdes 3.9.0 (options: --sc --careful --disable-rr) [39], and the contigs were qualified by QUAST 4.5 [40]. CheckM 1.0.6 [41] with the “lineage_wf’ command was used to assess the completeness and the contamination read rate after the contigs smaller than 1,000 bp were removed. Composite draft genomes were constructed by co-assembly of SAG groups with ≥99% identical marker genes detected by CheckM and ≥95% ANI. When an assembled composite draft genome showed ≥10% completeness, the composite draft was reassembled from the same SAG group by ccSAG [23]. CDS, rRNAs, and tRNAs were extracted from SAG contigs or composite draft genomes by prokka 1.13 [42], and the metabolic functions were estimated by Genomaple 2.3.2 [43]. Taxonomic annotation of draft genomes was performed by GTDB-Tk 0.3.0, [25], and *Bacteroides* phylogenetic trees were constructed by the neighbor-joining method using concatenated marker genes detected by CheckM. PUL clusters of *Bacteroides* composite genomes were detected by BLASTP [44] searches against the Bo cluster with default parameters, and phylogenetic trees were constructed from nucleotide sequences of the PUL clusters in the same manner. Orthologous groups were inferred by OrthoFinder 2.3.3 [45] for additional evaluation of genomic similarities among *Bacteroides* strains.

### Structure prediction

Homology modeling of proteins encoded by each gene included in the *Bacteroides* PUL cluster was performed using SWISS-MODEL [46] in automated mode. Based on the predicted protein structure, structural alignment was performed using MATRAS [47]. SusC-SusD complex structure construction was performed by MODELLER 9.20 [48] using template search and construction scripts created by HOMCOS [49] and using the crystal structure of BT1762-1763 (PDB_5T4Y) as a template.

## Additional files

**Additional file 1: Table S1.** Statistics for 346 SAGs obtained from the mouse fecal microbiome **Table S2.** Taxonomic annotation for 346 SAGs and 28 SAGCs obtained from the mouse fecal microbiome **Table S3.** Comparison of OTU detection between 16S rRNA gene amplicon sequencing and single-cell sequencing **Table S4.** Statistics for 28 composite SAGs (IMSAGCs) obtained from the mouse fecal microbiome. **Table S5.** Detailed statistics for medium and high-quality draft genomes obtained from the mouse fecal microbiome **Table S6.** Statistics for *Bacteroides* genomes used in comparative genome analysis **Table S7.** Numbers of signal peptide-containing predicted glycoside hydrolases/polysaccharide lyases **Table S8.** Statistics for high completeness SAGs (> 80%) associated with inulin uptake and TCA cycle-mediated metabolism **Table S9.** Statistics for conserved metabolic modules in qualified draft genomes and known *Bacteroides* spp. reference genomes

**Additional file 2: Figure S1.** Comparison of abundance profiles of IMSAGC_001 (OTU2) and IMSAGC_004 (OTU4) in 16S rRNA gene sequencing (a), (b) Violin plots indicate abundance profiles of IMSAGC_001 (OTU2) (a) and IMSAGC_004 (OTU4) (b) before and after two weeks of inulin or cellulose feeding in Lot 1 mice (n=5, Tukey’s HSD test). (c) Time-dependent changes in abundance profiles of IMSAGC_001 (OTU2) and IMSAGC_004 during 2-weeks inulin feeding in Lot 2 mice (n=9). **Figure S2** Predicted structures of sus homologs in IMSAGC_001 and IMSAGC_004 (a) Representation of the predicted SusC-SusD complex protein structure in IMSAGC_001. Units are shown in different colors. The crystal structure of BT1762-1763 (PDB_5T4Y) in *Bacteroides thetaiotaomicron* was used as template. (b), (c) Trees based on structural alignments of SusC (b) and SusD (c) among fructan-utilizing *Bacteroides* strains. **Figure S3.** Predicted partial reverse TCA cycles in IMSAGC 001 and IMSAGC 004 Succinate is formed by the reversal of partial TCA cycle reactions. Pyruvate is carboxylated to form oxaloacetate, which is then reduced to malate, fumarate, and succinate.

## Declarations

### Ethics approval and consent to participate

The protocols of all animal studies were approved by the Committee for Animal Experimentation of the School of Science and Engineering, Waseda University (No. 2017-A057 and 2018-A068)

### Consent for publication

Not applicable.

### Availability of data and materials

Sequence raw data obtained from mouse gut microbes and high- and medium quality draft genomes are listed in Table 1 and Additional file1: Data from Table S5 were deposited in the DNA Data Bank of Japan (DDBJ) under the accession number PRJDB8805.

### Authors’ contributions

M.H., R.C., and H.T. conceived and designed the experiments. M.H., Y.N., and K.M. developed the SAG-gel workflow. R.C. conducted animal experiments, metagenomic analysis, and metabolomic analysis. M.H., Y.N. M.K. C.S., and K.T. conducted single-cell genomics experiments and collected the data. M.H., R.C., M.K., and K.I conducted bioinformatic analysis of the metagenomic data and single-cell genomic data. M.H., R.C., M.K., K.I., and H.T. wrote the manuscript. All authors read and approved the final manuscript.

## Acknowledgments

The super-computing resource was provided by the Human Genome Center (University of Tokyo).

