## Additional file 2 for "Single-cell genomics of uncultured bacteria reveals dietary fiber responders in the mouse gut microbiota"

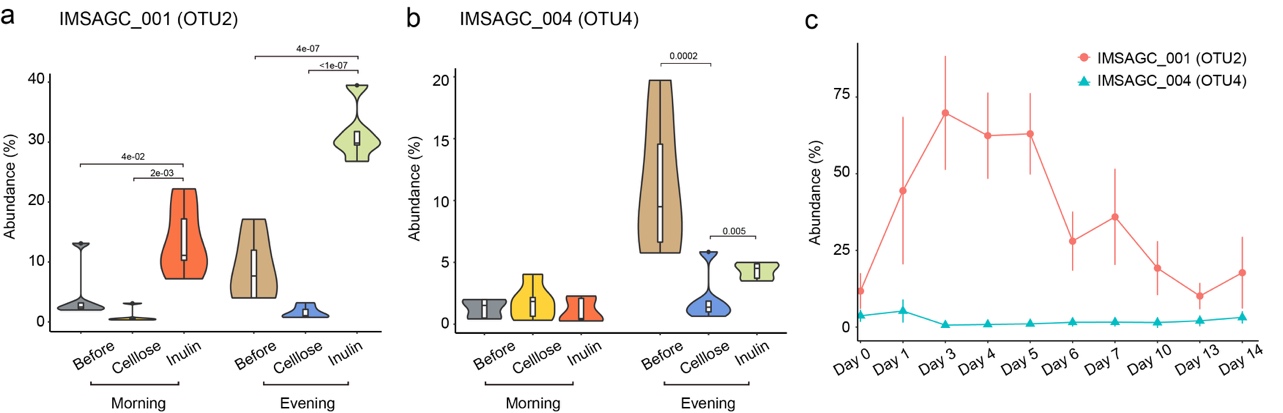

Figure S1. Comparison of abundance proﬁles of IMSAGC_001 (OTU2) and IMSAGC_004 (OTU4) in 16S rRNA gene sequencing (a), (b) Violin plots indicate abundance proﬁles of IMSAGC_001 (OTU2) (a) and IMSAGC_004 (OTU4) (b) before and after two weeks of inulin or cellulose feeding in Lot 1 mice (n=5, Tukey's HSD test). (c) Time-dependent changes in abundance proﬁles of IMSAGC_001 (OTU2) and IMSAGC_004 during 2-weeks inulin feeding in Lot 2 mice (n=9).

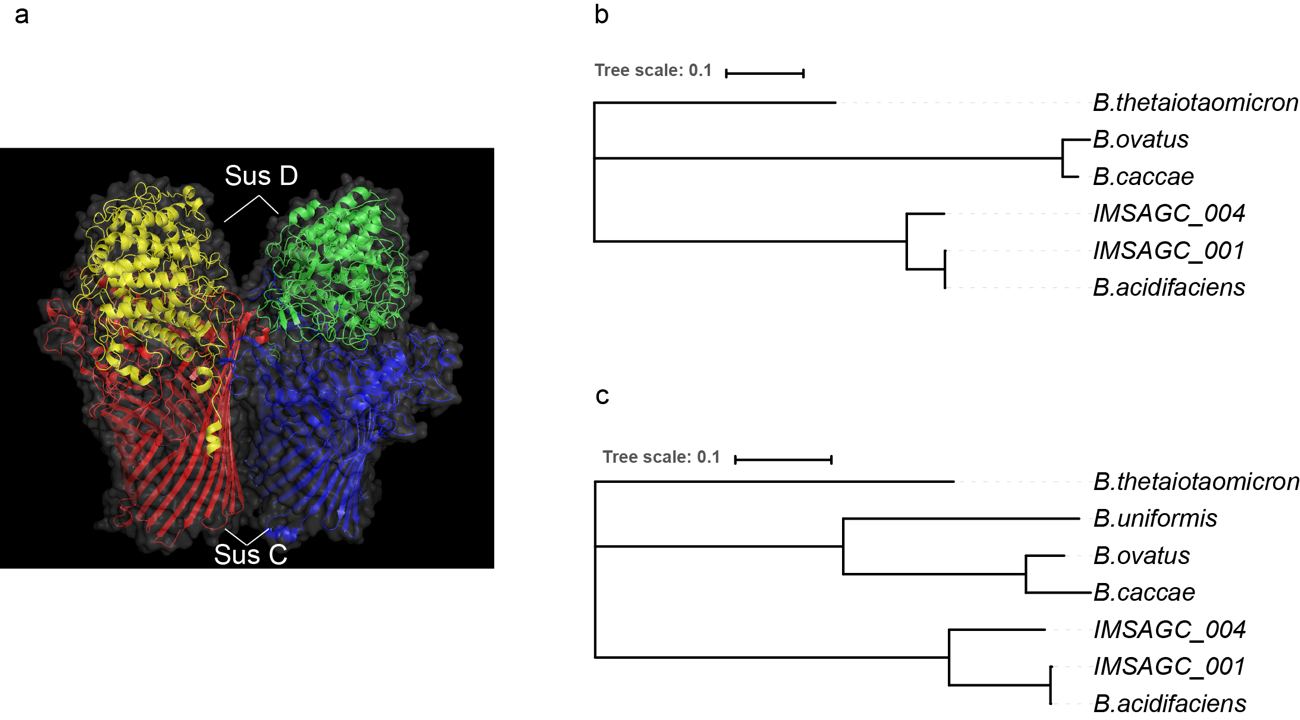

Figure S2. Predicted structures of sus homologs in IMSAGC_001 and IMSAGC_004 (a) Representation of the predicted SusC-SusD complex protein structure in IMSAGC_001. Units are shown in different colors. The crystal structure of BT1762-1763 (PDB_5T4Y) in *Bacteroides thetaiotaomicron* was used as template. (b), (c) Trees based on structural alignments of SusC (b) and SusD (c) among fructan-utilizing *Bacteroides* strains.

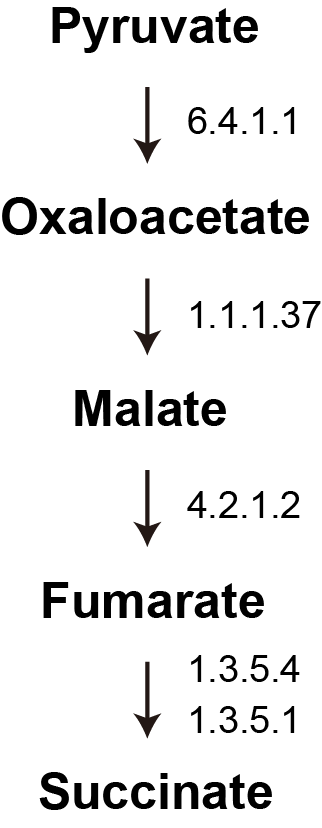

Figure S3. Predicted partial reverse TCA cycles in IMSAGC 001 and IMSAGC 004 Succinate is formed by the reversal of partial TCA cycle reactions. Pyruvate is carboxylated to form oxaloacetate, which is then reduced to malate, fumarate, and succinate.
